# The hippocampus contributes to retroactive stimulus associations during trace fear conditioning

**DOI:** 10.1101/2022.10.17.512614

**Authors:** Kyle Puhger, Ana P. Crestani, Cassiano R. A. F. Diniz, Brian J. Wiltgen

## Abstract

Binding events that occur at different times is essential for memory formation. In trace fear conditioning, animals associate a tone and footshock despite no temporal overlap. The hippocampus is thought to mediate this learning by maintaining a memory of the tone until shock occurrence, however evidence for sustained hippocampal tone representations is lacking. Here we demonstrate a retrospective role for the hippocampus in trace fear conditioning. Bulk calcium imaging revealed sustained increases in CA1 activity after footshock that were not observed after tone termination. Optogenetic silencing of CA1 immediately after footshock impaired subsequent memory. Additionally, footshock increased the number of sharp wave-ripples compared to baseline during conditioning. Therefore, post-shock hippocampal activity likely supports learning by reactivating and linking latent tone and shock representations. These findings highlight an underappreciated function of post-trial hippocampal activity in enabling retroactive temporal associations during new learning, as opposed to persistent maintenance of stimulus representations.

## Introduction

A major function of the hippocampus is to learn about events that are separated in space and time^1–5^. This information is used to form episodic memories in humans and spatial maps in animals. In the latter case, proximal and distal stimuli are associated during high-frequency oscillations called sharp-wave ripples (SWRs)^6–13^. For example, when a rat discovers food at the end of a maze CA1 neurons replay (or reactivate) the path that was taken to get there^6–9,13,14^. These replay events occur rapidly (150-300ms) and coincide with the release of neuromodulators like dopamine (DA), which strengthens the connections between co-active neurons so the entire path will be remembered in the future^9,15^.

It is possible that a similar process allows animals to learn about non-spatial events that are separated in time. For example, during trace fear conditioning (TFC), an aversive footshock (US) is presented several seconds after the termination of an auditory cue (CS). Animals can associate these stimuli provided they have an intact hippocampus^16–20^. However, it is not known if SWRs or replay contribute to this learning. It is typically assumed that the hippocampus maintains a memory of the auditory cue after it terminates so it can be associated with shock several seconds later ^20–23^. Although this idea is widely accepted, recording and imaging studies have observed little to no maintenance of “CS activity” after the auditory cue has terminated ^24,25^. Instead, the hippocampus is transiently activated during the CS and then again during and after footshock. Interestingly, the activity induced by footshock is prolonged and persists for tens of seconds after this stimulus terminates^24,25^. We hypothesize that this period of prolonged activity represents a period of time when memories of the tone and shock are reactivated together and become associated, similar to that observed during spatial learning.

The idea that learning occurs after a conditioning trial has ended was first proposed by animal researchers in the 1940s, 50s and 60s ^26–29^. Edward Tolman wrote, “*Learning what to avoid…may occur exclusively during the period after the shock.”.* Leo Kamin hypothesized that unexpected shocks cause animals to perform, “*…a backward scanning of their memory store of recent stimulus input; only as a result of such a scan can an association between CS and US be formed.”.* In the 1970s, Allen Wagner reintroduced this idea and argued that, “*an unexpected or surprising US event will provoke a post-trial rehearsal process necessary to the learning about the CS-US relationships involved.*”^30^. The assertion that surprise is essential for learning later became widely accepted in psychology and drove the development of many influential learning theories^31,32^. However, the assumption that unexpected stimuli induce memory formation because they instigate a post-trial rehearsal process has largely disappeared from the field. This is despite the fact that a number of behavioral studies provided support for this idea^28–30,33–36^.

In the current paper, we test the idea that hippocampal activity after the US is essential for trace fear learning. We hypothesize that this activity consists of SWRs, during which, memories of the CS and US are reactivated and become associated with one another. The post-trial period is ideal for learning because it is when neuromodulators like dopamine (DA) are released into the hippocampus^37^. DA is well-known to enhance synaptic plasticity and promote long-term memory formation^38–45^. Consistent with this idea, we recently showed that DA activity is required for TFC and levels of this catecholamine increase in dorsal CA1 for approximately 40-sec after footshock^37^. To test our current hypotheses, we used a combination of bulk calcium-imaging, optogenetics, *in vivo* recordings and pharmacological manipulations during trace fear learning in mice.

## Results

### Footshock elicits a large increase in CA1 calcium activity

We used fiber photometry to measure bulk calcium fluorescence, an indirect readout of population activity, in order to examine neural activity in CA1 during TFC. Mice were injected with CaMKII-GCaMP6f which expresses the calcium indicator GCaMP6f in CA1 pyramidal cells (Fig. 1A). Mice then underwent a single session of TFC training consisting of 10 training trials. Consistent with previous studies we did not find any significant GCaMP response to the CS onset, CS offset, or during the trace interval (Fig. 1B)^24,25^. However, we found a large, sustained increase in GCaMP fluorescence elicited by the US (Fig. 1B). The mean GCaMP fluorescence was significantly greater during the 20 s after the footshock (post-shock) than during the 20 s prior to the shock (pre-shock) (Fig 1C). These data demonstrate a large US-elicited increase in CA1 activity, raising the possibility that this US-induced activity also contributes to TFC learning. Next, we will test this hypothesis by using optogenetic inhibition to selectively silence CA1 after the footshock during TFC. We predict that silencing CA1 immediately after the footshock, but not later during the ITI, will impair memory.

**Figure 1.**
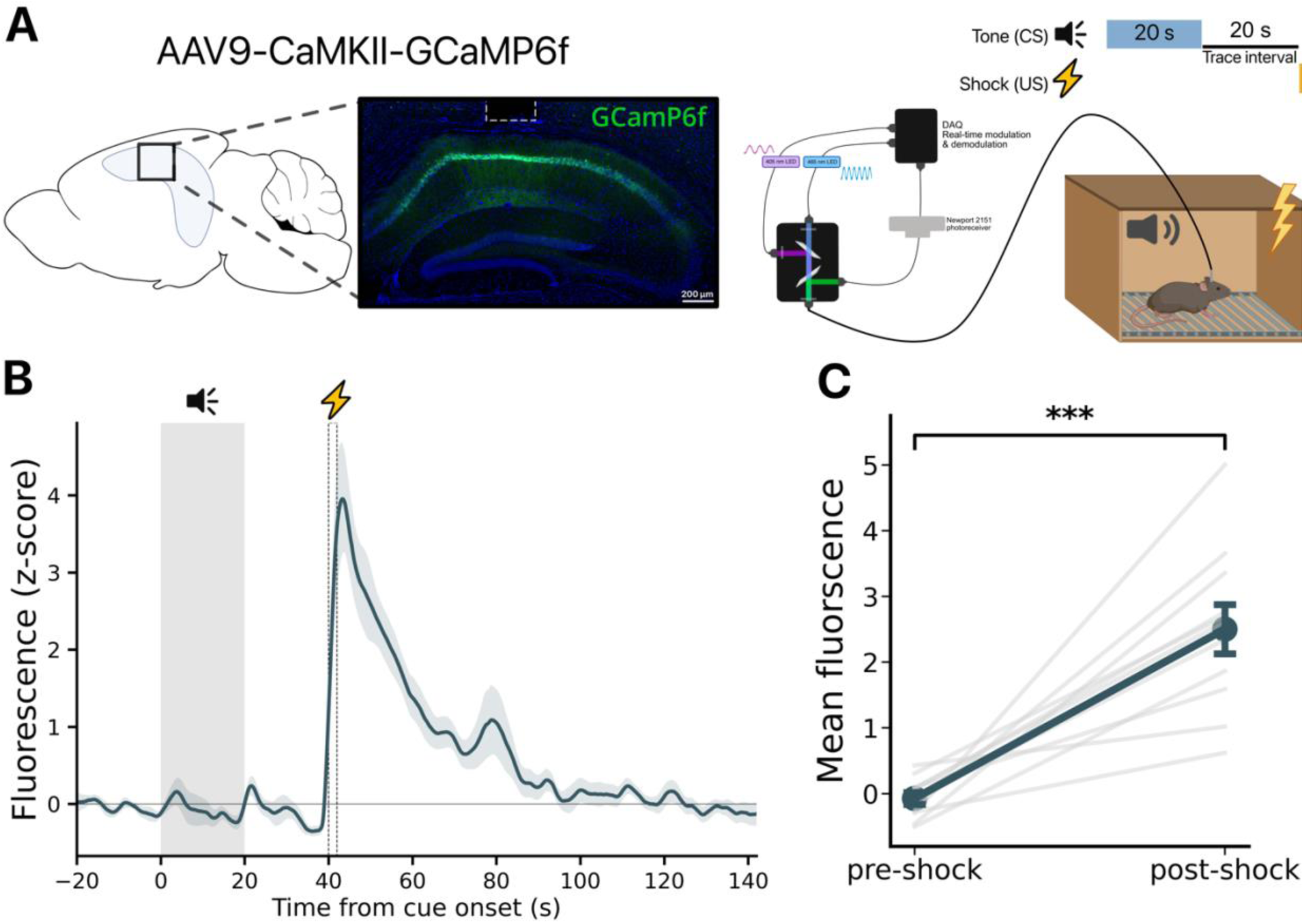
Footshock elicits increases in CA1 activity during TFC training. **(A)** *Left:* GCaMP6f was infused into CA1 (n = 11 mice), and an optical fiber (white dotted line) was implanted above the infusion site. Right: Schematic of the fiber photometry system used to measure bulk fluorescence during TFC training. **(B)** Bulk calcium response during TFC training trials show a large increase in activity elicited by the footshock. Gray rectangle indicates when tone is presented. Dotted rectangle indicates footshock presentation. **(C)** GCaMP fluorescence is significantly increased by footshock (paired t-test, t(10) = −6.256, p < 0.001). Light gray lines represent each animal’s mean fluorescence for the 20 seconds before the shock (pre-shock) and the 20 seconds after the shock (post-shock). Dark line represents the mean pre-shock and post-shock fluorescence averaged over all subjects. All data are expressed as mean ± SEM; *p < 0.05, **p < 0.01, ***p < 0.001.

### CA1 inhibition after the footshock impairs TFC memory

After observing a large increase in US-elicited CA1 pyramidal cell activity, we next sought to determine whether CA1 activity during the post-shock period was necessary for TFC learning. To test this we, infused either AAV-CaMKII-eArchT3.0-eYFP to express the inhibitory opsin ArchT or AAV-CaMKII-eGFP in CA1 pyramidal cells. During training, 561 nm light was delivered continuously for 40 s immediately after the footshock for all three CS-US pairings (Figure 2B,C). During training, there were no group differences in freezing during the baseline period prior to conditioning, but ArchT mice froze significantly less than eGFP mice during the trace interval and ITI (Figure 2D). When tone memory was tested the next day in a novel context in the absence of laser stimulation, ArchT mice froze significantly less than eGFP controls (Figure 2E). Twenty-four hours later we assessed context memory by returning the mice to the training context for 600 s. ArchT mice froze significantly less than eGFP controls (Figure 2F). A deficit in context fear was not observed previously when CA1 was inactivated during the tone or trace interval^16,18^. This suggests that CA1 activity immediately after footshock supports memory for both the tone and context in TFC.

**Figure 2.**
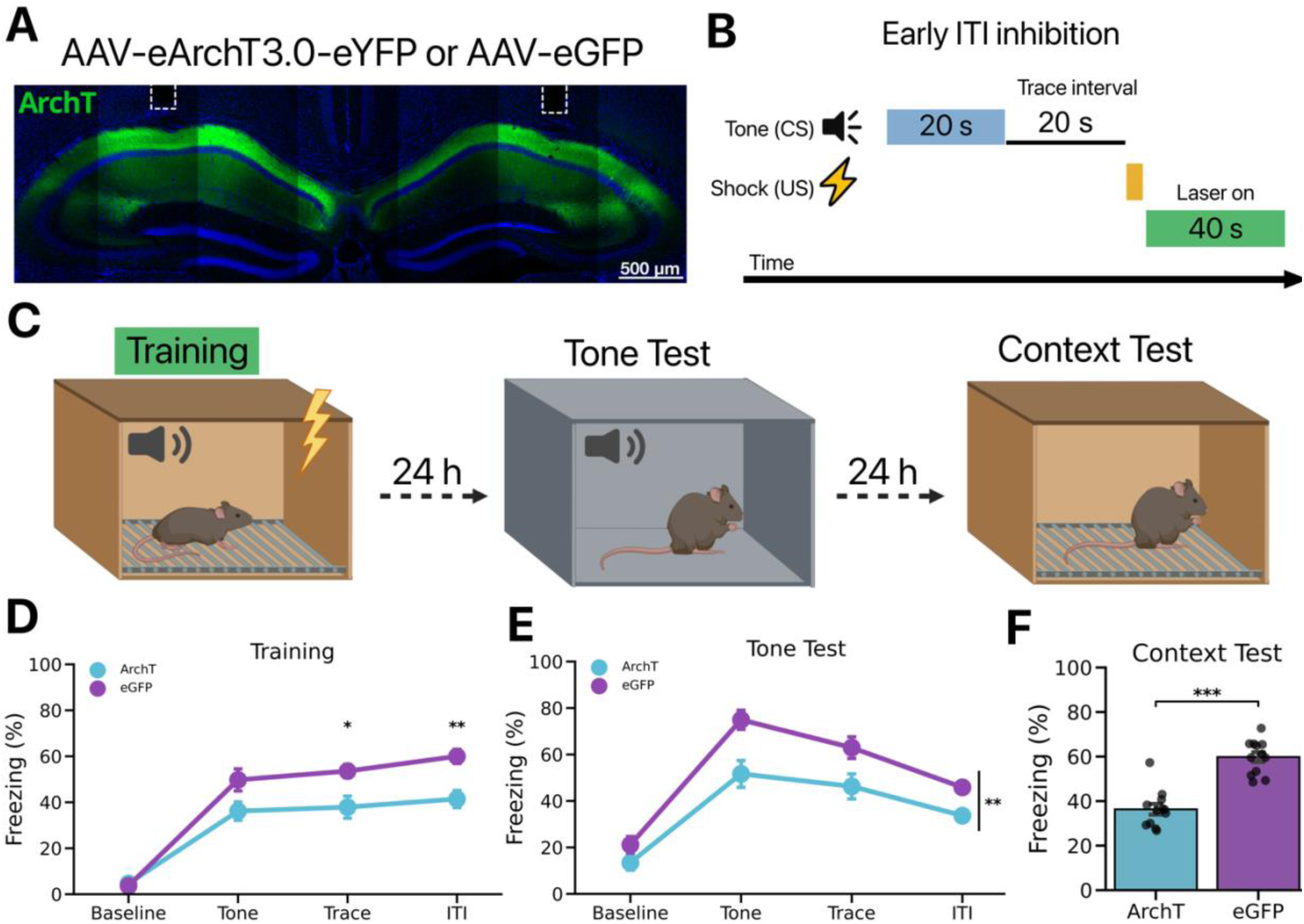
CA1 inhibition after the footshock impairs TFC memory. **(A)** Representative image of AAV expression and optical fiber placement targeting CA1. **(B)** Experimental design to silence CA1 after the footshock during TFC. **(C)** On the first day mice (n = 12 per group) underwent TFC while laser stimulation (561 nm) was delivered to CA1 continuously for 40 s immediately after the footshock on each training trial. The next day mice received a tone test in a novel context B. The following day contextual fear memory was tested in the original training context. **(D)** ArchT mice froze significantly less during the trace interval and ITI than the eGFP control group during the training session (Group x Phase interaction: F(3,66) = 4.842, p < 0.01; Trace: t(22) = −2.801, p < 0.05; ITI: t(22) = −3.887, p < .01). **(E)** ArchT mice froze significantly less than eGFP mice during the tone test (Main effect of Group: F(1,22) = 11.32, p < 0.01). **(F)** ArchT mice froze significantly less than eGFP mice during the context test (t(22) = −7.17, p < 0.001). All data are expressed as mean ± SEM; *p < 0.05, **p < 0.01, ***p < 0.001.

### Delayed CA1 inhibition during the ITI does not impair TFC memory

Prior work has demonstrated that CA1 activity during the trace interval is critical for TFC memory^16–18^. Our current results indicate that CA1 activity after the footshock is also necessary for TFC memory formation (Figure 2B–C). In order to rule out any potential nonspecific effects of CA1 inhibition during training, we repeated the previous optogenetic inhibition experiment but delayed inhibition until later in the ITI. Mice received injections of ArchT or eGFP into dorsal CA1 (n = 12 per group). Three weeks later mice were trained as described in the previous experiment, but laser stimulation was presented 140 s after each footshock (Figure 3B,C). Unlike the results from the early ITI inhibition, there were no differences in freezing between ArchT and eGFP mice during training (Figure 3D). Additionally, when tone memory was tested the following day in a novel context both groups of mice displayed similar levels of freezing to the tone (Figure 3E). This is consistent with previous findings that CA1 disruption during the ITI does not affect tone memory in TFC^17,18^. Similarly, context memory was also unaffected when CA1 inhibition after the footshock was delayed (Figure 3F). These results provide evidence that CA1 is selectively required immediately after the footshock but not later in the ITI.

**Figure 3.**
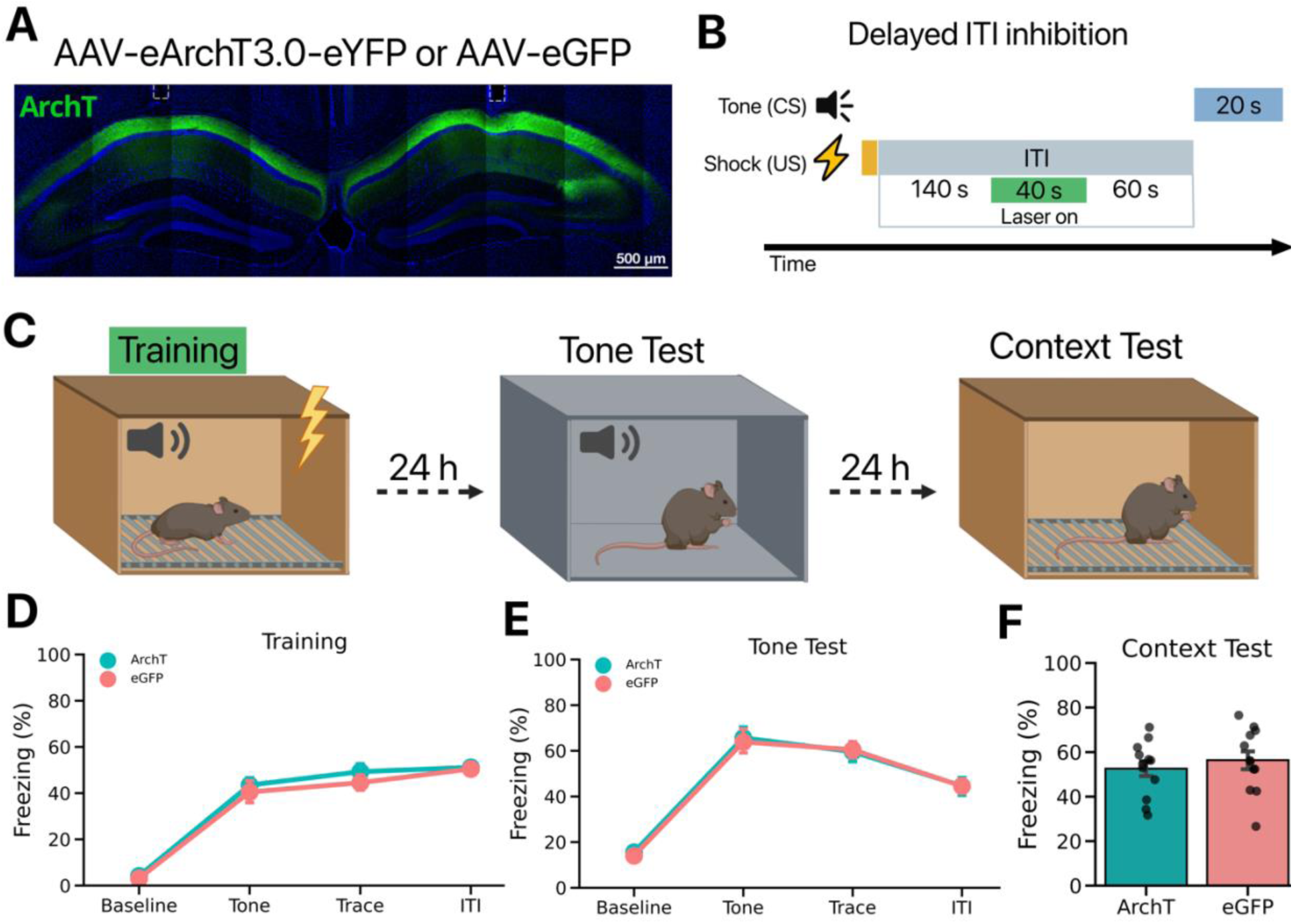
Delayed CA1 inhibition does not impair TFC memory. **(A)** Representative image of post hoc validation of AAV expression and optical fiber placement (white dotted lines) targeting CA1. **(B)** Experimental design to silence CA1 during the ITI. On the first day mice underwent TFC while laser stimulation (561 nm) was delivered to CA1 continuously for 40 s starting 140 s after the footshock on each training trial. **(C)** On the first day mice underwent TFC while laser stimulation (561 nm) was delivered to CA1 continuously for 40 s after a 140 s delay following termination of the footshock. The next day mice received a tone test in a novel context B. The following day contextual fear memory was tested in the original training context. **(D)** ArchT mice and eGFP performed similarly during training (Main effect of Group: F(1,22) = 0.446, p > 0.05). **(E)** ArchT mice and eGFP did not differ in their freezing to the tone CS during the tone test (Main effect of Group: F(1,22) = 0.022, p > 0.05). **(F)** Both groups showed similar freezing responses to the training context During the contextual memory test (t(22) = −0.694, p > 0.05). All data are expressed as mean ± SEM; *p < 0.05, **p < 0.01, ***p < 0.001.

### Post-shock CA1 inhibition late in learning does not impair TFC memory

Our results thus far suggest that CA1 may contribute to TFC learning by retroactively associating the US and CS. Next, we asked whether CA1 activity after the footshock was involved in the maintenance of previously consolidated memories. To test this idea, we injected mice with ArchT or eGFP (n = 12 per group) as described in the previous experiments. On the first day, mice were given 3 TFC trials in the without laser stimulation (Figure 4A). No group differences were observed during this session (Figure 4B). On training day 2 mice were given another 3 TFC trials, and laser stimulation was delivered 40 s immediately after the footshock (Figure 4A). Contrary to post-shock CA1 inactivation during initial learning, silencing CA1 on the second day of training did not impair learning (Figure 4C). During the tone test both groups of mice froze similarly in response to the tone (Figure 4D). Contextual fear memory was also similar between groups (Figure 4E). These results indicate that silencing CA1 immediately after the footshock do not impair a previously formed TFC memory. This is consistent with the view that CA1 activity after the footshock is required to initially learn the CS-US relationship but is not required when animals have already learned the CS-US association.

**Figure 4.**
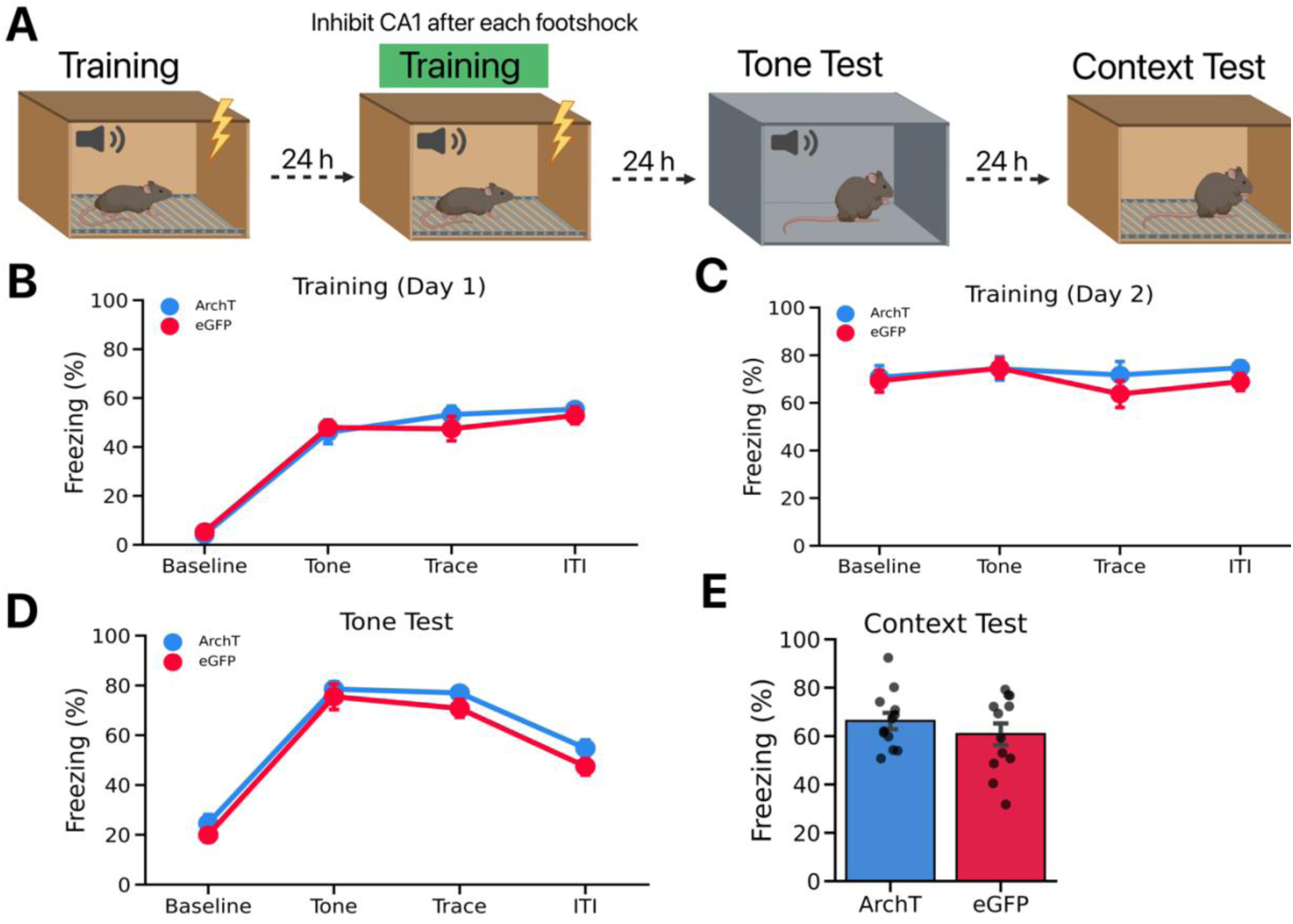
Post-shock CA1 inhibition late in learning does not impair TFC memory. **(A)** Experimental design to silence CA1 after the footshock on the second training day. Both groups received TFC training on the first day without any laser stimulation. On the second training day, mice underwent TFC while laser stimulation (561 nm) was delivered to CA1 continuously for 40 s immediately after the footshock on each training trial. The next day mice received a tone test in a novel context B. The following day contextual fear memory was tested in the original training context. **(B)** ArchT mice and eGFP performed similarly during the first day of training (Main effect of Group: F(1,22) = 0.147, p > 0.05). **(C)** ArchT mice and eGFP performed similarly during the second training day (Main effect of Group: F(1,22) = 0.690, p > 0.05). **(D)** During the tone test, ArchT mice and eGFP did not differ in their freezing to the tone (Main effect of Group: F(1,22) = 1.88, p > 0.05). **(E)** During the context test, both groups showed similar freezing responses to the training context (t(22) = 0.944, p > 0.05). All data are expressed as mean ± SEM; *p < 0.05, **p < 0.01, ***p < 0.001.

### Footshock during TFC produces an increase in the occurrence of SWRs

To test whether SWRs are increased after the footshock, we conducted electrophysiological recording of CA1 LFPs (local field potential) while mice underwent a single session of TFC training, consisting of 10 tone-trace-shock trials (Figure 5A,B). We found that the number of SWRs was significantly greater after the footshock than during the baseline period (Figure 5C). These data indicate that shock elicits an increase in SWR occurrence, which we hypothesize contributes to retroactive learning in TFC.

**Figure 5.**
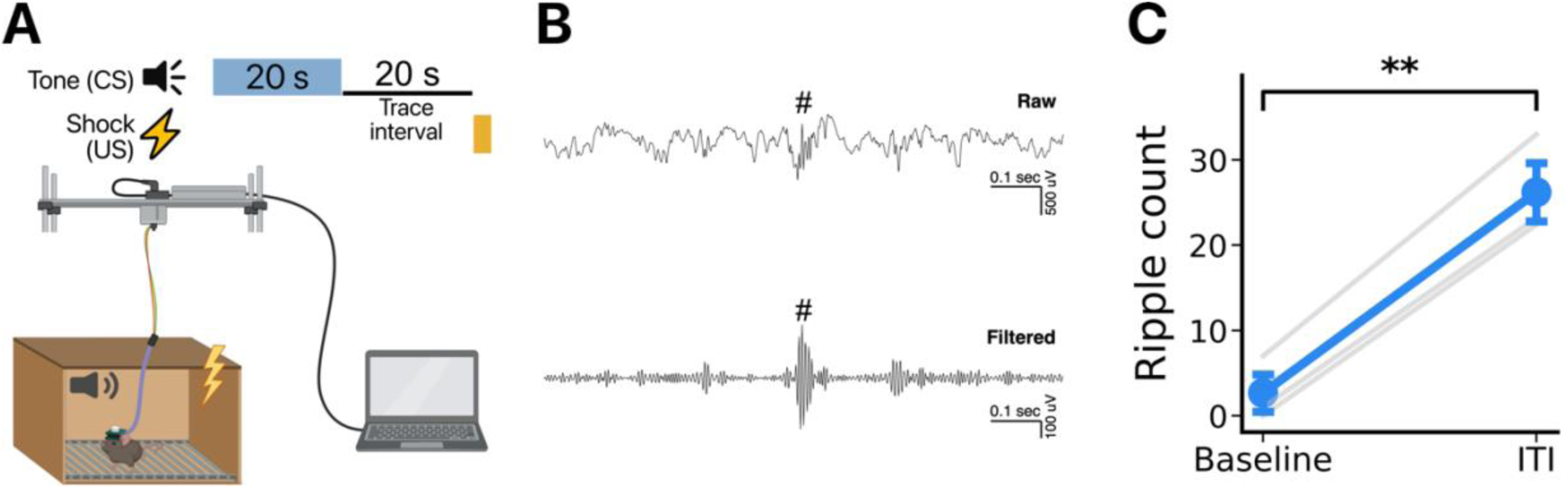
SWRs increase in frequency after the US. **(A)** Schematic of the electrophysiological setup used to record local field potentials (LFPs) during TFC training. **(B)** Representative LFP recording. Top: raw data example of LFPs. Bottom: 130-200Hz filtered data. Octothorpe (#) indicates detected SWR. **(C)** SWR frequency is significantly increased after the shock (t(2) = −19.06, p < 0.01). All data are expressed as mean ± SEM; *p < 0.05, **p < 0.01, ***p < 0.001.

### Muscarinic cholinergic agonist decreases SWRs, reduces calcium activity during TFC and impairs memory

Prior work has demonstrated that increased acetylcholine suppresses ripples^47^. Similarly, others have shown that pilocarpine, a muscarinic cholinergic agonist, decreases the frequency of SWRs^48^. Here, we first demonstrated that an I.P. injection of pilocarpine (10 mg/kg) reduced the frequency of SWRs ≈ 3x in anesthetized animals (Figure 6A). To determine the effects of acetylcholine on bulk calcium activity and TFC memory, we infused GCaMP6f into CA1 and subsequently administered either pilocarpine or saline (n = 5 mice per group) 15 mins prior to TFC training (Figure 6B). We found that GCaMP fluorescence was significantly lower in the pilocarpine group compared with the saline group in the 20 s period after footshock (Figure 6C). Additionally, pilocarpine injected mice showed a memory impairment in the retrieval session performed 24h after training (Figure 6D). Together, these results suggest that the increase in cholinergic transmission induced by pilocarpine decreases the frequency of SWRs, reduces CA1 activity after footshock and impairs memory.

**Figure 6.**
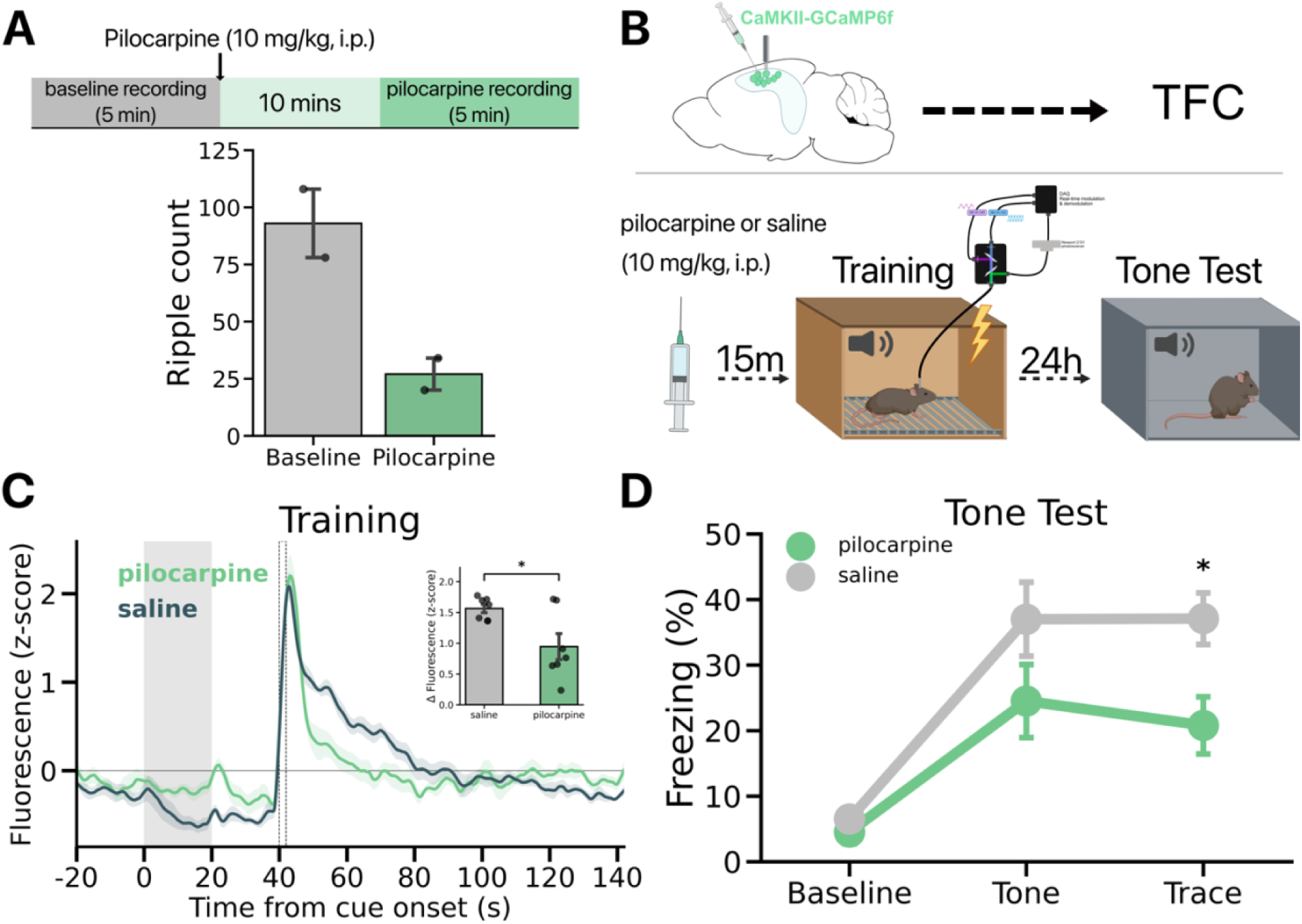
Effects of cholinergic agonism on SWRs, GCaMP activity, and TFC memory. **(A)** Mice (n = 2) were first anesthetized prior to a 5 mins baseline LFP recording (baseline) in their home cage prior to receiving an injection of pilocarpine (10 mg/kg, i.p.). After a 10 min delay, mice underwent another 5 mins of LFP recording (pilocarpine). **(B)** Schematic of TFC design. GCaMP was infused into the dorsal hippocampus and an optical fiber was implanted above CA1 prior to mice undergoing TFC. Saline or pilocarpine (10 mg/kg, i.p.) was administered 15 mins prior to TFC training (n = 5 per group) wherein bulk calcium activity was measured by fiber photometry. The next day, mice underwent a tone test in a novel context. **(C)** Trial-averaged GCaMP fluorescence in saline and pilocarpine mice. *Inset:* GCaMP fluorescence was significantly lower after the footshock in pilocarpine mice compared to saline mice (post – pre shock: t(12) = −2.787, p = 0.016). **(D)** Mice that received pilocarpine injections prior to training froze significantly during the trace interval than mice that received saline prior to training (Group x Phase interaction: F(2,24) = 3.579, p < 0.05; Trace t(12) = −2.771, p < 0.05). All data are expressed as mean ± SEM; *p < 0.05, **p < 0.01, ***p < 0.001.

## Discussion

Trace conditioning depends on the hippocampus or its homologue in a variety of specifies including fish, rodents, rabbits and humans^19,20,49–53^. This is likely the case because it engages a fundamental function of this brain region; associating stimuli that are separated in time and space^1–5^. Consistent with this idea, when animals learn to associate stimuli that overlap in time (e.g. delay conditioning), the hippocampus is not required^17,19,54–56^. This suggests that other brain regions can link stimuli provided they receive input about them very close together in time. We hypothesize that this unique function of the hippocampus underlies the formation of episodic memories in humans and spatial/cognitive maps in animals^3,4^.

One way the hippocampus associates discontiguous events is via SWRs and replay. For example, in spatial learning tasks, animals are trained to navigate a maze in order to receive food reward. As they travel towards the food, place cells fire at different spatial locations on the maze. When food is obtained, the same place cells are reactivated in reverse order during SWRs from the end of the maze back to the beginning^6–8^. This occurs very quickly (150-300ms), which is thought to strengthen the connections between place cells so the path can be remembered in the future.

We hypothesize a similar process takes place during TFC. That is, when animals receive an aversive shock, SWRs are induced and the hippocampus replays the sequence of events that preceded it. Previous work has shown that extended replay events can stretch backwards in time for at least 30-40 sec^13,57,58^. If that were to happen during TFC, memory for the CS (distal cue) would be reactivated at the same time as memory for the shock (proximal cue). However, this does not imply that the hippocampus encodes the emotional value of shock like the amygdala and other brain regions^59^. Instead, it is thought to form a sensory representation of this event like it does for reward and other cues that are encountered in the environment^60,61^.

To test our hypothesis, we first used fiber photometry to record bulk calcium fluorescence from CA1 during TFC. We found no change in bulk calcium fluorescence in response to the CS or during the trace interval. The lack of a CS-evoked response is likely due to the small number of CA1 neurons that respond to pure tones^62–64^. The use of more complex auditory stimuli should make this response easier to observe in future studies^65^. The absence of a change in CA1 activity during the trace interval is consistent with a large number of recording and imaging studies, as noted above^24,25,66,67^. This result casts doubt on the idea that CA1 neurons maintain an active memory trace of the CS after it terminates.

In contrast to the auditory cue and trace interval, our fiber photometry recordings showed that footshock elicits a large and prolonged increase in CA1 activity. These data are congruent with prior work showing that pyramidal cells in CA1 are strongly activated by aversive unconditional stimuli^24,25^. We next sought to determine whether the increased post-shock activity in CA1 was causally involved in the acquisition of TFC memory. First, we found that optogenetic inhibition of CA1 during the period when activity is elevated by footshock led to a significant memory impairment for both the tone and training context. This result is similar to that observed in eyeblink conditioning studies where surprising or unexpected post-trial events were shown to interfere with learning, presumably by disrupting a post-trial “rehearsal” process^30,36^. Based on these data, we hypothesize that CA1 plays a role in the retroactive processing of the CS-US relationship.

We also found that delaying CA1 inactivation until 140 s after the footshock did not impair TFC memory. This is consistent with previous studies showing that CA1 inactivation during the ITI does not affect trace fear acquisition^17,18^. We also found that post shock activity is most important early in learning when US prediction errors and dopamine release are largest in CA1^37^. When post-shock inactivation occurred during a second training session (24 hours after initial learning) there was no effect on memory. These results suggest that CS-US associations are formed when unexpected shocks increase CA1 activity and induce dopamine release in the hippocampus.

When place cells were discovered, O’Keefe and Nadel argued that the main function of the hippocampus was to generate a map of the environment that animals could use to guide subsequent behavior^68^. Subsequent recording studies demonstrated that the hippocampus binds spatial and non-spatial information together to generate an internal model of the world (i.e. a “cognitive map”) wherein space is only one of several relevant dimensions^4,26,68–70^. For example, the hippocampus has been shown to encode non-spatial information such as odors^71^, sound frequencies^64^, temporal intervals^72–75^ and abstract task variables when it is relevant to an animal’s behavior^76–79^. These data support the idea that the hippocampus forms associations between salient internal and external stimuli that then become stored in long-term memory^60^.

The hippocampus reactivates or replays sequences of activity during large bursts of population activity known as sharp wave-ripples (SWRs)^9,10^. Importantly, these events can be replayed in forward or reverse order which could support prospective and retrospective associations during behavior. Although hippocampal replay is often studied in the context of spatial behaviors, recent work has extended these findings to non-spatial tasks. For example, in a sensory preconditioning task it was found that neurons in CA1 representing reward outcome fired before neurons that represented the sensory cue during SWRs^80^.

It is important to note that reward and aversion are not required to form cognitive maps^26,81^. Replay occurs automatically after an animal explores and then pauses, becomes inactive or goes to sleep^10^. However, rewarding and aversive stimuli strengthen spatial memories by inducing the release of neuromodulators like dopamine and norepinephrine^42–44^. In addition, reward had been shown to promote replay of recently experienced events as opposed to future events^82^. These results suggest that spatial/cognitive maps are formed automatically, and their stability is enhanced by biologically relevant events such as food or pain.

SWRs are thought to coordinate activity throughout the brain. We hypothesize that US-induced increases in CA1 activity are driven, in part, by SWRs, which facilitate communication between the hippocampus and other brain areas like the amygdala^83^. It is possible, therefore, that SWRs transmit information about recently encountered stimuli to the amygdala (like the CS), which allows them to become associated with fear. The precise timing of this signal may not be important, as amygdala activity remains elevated for several seconds after an aversive event occurs^84–87^. Consequently, the convergence of SWRs with elevated amygdala activity could promote synaptic strengthening and allow memory representations in the hippocampus to drive defensive behaviors like freezing. Future work in our lab will focus on investigating single-unit activity in the hippocampus and amygdala during TFC. These data should reveal how interactions between these regions during the post-shock period support retroactive learning of the CS-US relationship.

## Methods

### Subjects

Subjects in this study were 8–16-week-old male and female mice (C57BL/6J, Jackson Labs; B6129F1, Taconic). Mice were maintained on a 12h light/12h dark cycle with *ad libitum* access to food and water. All experiments were performed during the light portion of the light/dark cycle (0700-1900). Mice were group housed throughout the duration of the experiment. All experiments were reviewed and approved by the UC Davis Institutional Animal Care and Use Committee (IACUC).

### Surgery

Stereotaxic surgery was performed 2-3 weeks before behavioral experiments began. Mice were anesthetized with isoflurane (5% induction, 2% maintenance) and placed into a stereotaxic frame (Kopf Instruments). An incision was made in the scalp and the skull was adjusted to place bregma and lambda in the same horizontal plane. Small craniotomies were made above the desired injection site in each hemisphere. AAV was delivered at a rate of 2nl/s to dorsal CA1 (AP - 2.0 mm and ML ± 1.5 mm from bregma; DV −1.25 mm from dura) through a glass pipette using a microsyringe pump (UMP3, World Precision Instruments). For the optogenetic inhibition experiments, the constructs were AAV5-CaMKIIa-eArchT3.0-EYFP (250 nl/hemisphere, titer: 4 × 10^12^, diluted 1:10, UNC Vector Core) and AAV5-CaMKIIa-GFP (250 nl/hemisphere, titer: 5.3 × 10^12^, diluted 1:10, UNC Vector Core). After AAV infusions, an optical fiber (optogenetics: 200 µm diameter, RWD Life Science, fiber photometry: 400 µm diameter, Thorlabs) was implanted above dorsal CA1 (AP −2.0 mm and ML ± 1.5 mm from bregma; DV −1.0 mm from dura). The fiber implants were secured to the skull using dental adhesive (C&B Metabond, Parkell) and dental acrylic (Bosworth Company). Optogenetic inhibition and fiber photometry recordings took place ∼2-3 weeks after surgery.

For the microdrive implantation, a ∼ 2 × 2 mm craniotomy was performed above the right hemisphere of the CA1 subregion of the dorsal hippocampus. One stainless-steel screw fixed in the interparietal bone area on the contralateral site served as ground reference for the recordings. The base of the microdrive and the screw were secured to the skull using dental adhesive (C&B Metabond, Parkell) and dental acrylic (Bosworth Company). The microdrive then was covered by a cap to avoid any damage.

### Behavioral apparatus

The behavioral apparatus has been described previously^16^. Briefly, fear conditioning occurred in a conditioning chamber (30.5 cm × 24.1 cm × 21.0 cm) within a sound-attenuating box (Med Associates). The chamber consists of a front-mounted scanning charge-coupled device video camera, stainless steel grid floor, a stainless-steel drop pan, and overhead LED lighting capable of providing broad spectrum and infrared light. For context A, the conditioning chamber was lit with both broad spectrum and infrared light and scented with 70% ethanol. For context B, a smooth white plastic insert was placed over the grid floor and a curved white wall was inserted into the chamber. Additionally, the room lights were changed to red light, only infrared lighting was present in the conditioning chamber, and the chamber was cleaned and scented with disinfectant wipes (PDI Sani-Cloth Plus). In both contexts, background noise (65 dB) was generated with a fan in the chamber and HEPA filter in the room.

### Trace fear conditioning

All behavioral experiments took place during the light phase of the light-dark cycle. Prior to the start of each experiment, mice were habituated to handling and tethering to the optical fiber patch cable for 5 mins/day for 5 days. Next, mice underwent training in context A. For optogenetic inhibition experiments, mice were allowed to explore the conditioning chamber during training for 240 s before receiving three conditioning trials. Each trial consisted of a 20-second pure tone (85 dB, 3 kHz), a 20 s stimulus-free trace interval, and a 2 s footshock (0.4 mA) followed by an intertrial interval (ITI) of 240 s. The following day, mice were placed in a novel context (context B) for a tone memory test consisting of a 240 s baseline period followed by six CS presentations separated by a 240 s ITI. Twenty-four hours later mice were returned to the training context A for 600 s to test their context memory. For fiber photometry experiments, mice were allowed to explore the conditioning chamber during training for 120 s before receiving ten conditioning trials. Each trial consisted of a 20-second pure tone (85 dB, 3 kHz), a 20-second stimulus-free trace interval, and a 2-second footshock (0.3 mA) followed by an intertrial interval (ITI) of 120 s. Freezing behavior was measured using VideoFreeze software (Med Associates) and processed using custom python scripts.

### Optogenetic inhibition of CA1

For optogenetic inhibition experiments green light (561 nm, ∼10 mW) was delivered continuously for 40 s during each training trial. No light was delivered during the tone or context memory tests. For both post-shock silencing experiments light was delivered immediately after termination of the footshock. For the ITI silencing experiment light was delivered 140 s after termination of the footshock.

### Fiber photometry

Fiber photometry enables the measurement of bulk fluorescence signal from a genetically defined population of cells in freely-moving, behaving mice. To characterize bulk CA1 pyramidal cell bulk calcium activity, we expressed GCaMP6f under the CaMKII promoter and a 400 µm 0.37 NA low autofluorescence optical fiber was implanted above the injection site. The fiber photometry system (Doric) consisted of an FPGA based data acquisition system (Fiber Photometry Console, Doric) and a programmable 2-channel LED Driver (Doric) to control two connectorized light-emitting diodes (LED): a 465 nm LED (to measure calcium-dependent changes in GCaMP fluorescence) and a 405 nm LED (an isosbestic control channel that measures calcium-independent changes in fluorescence). LED power was set to ∼40 µW, and the LEDs were modulated sinusoidally (465 nm at 209 Hz, 405 nm at 311 Hz) to allow for lock-in demodulation of the source signals. Light was passed through a sequence of dichroic filters (Fluorescent Mini Cube, Doric) and transmitted into the brain via the implanted optical fiber. Bulk GCaMP fluorescence from pyramidal cells beneath the optical fiber was collected and passed through a GFP emission filter (500-540 nm) and collected on a femtowatt photoreceiver (Newport 2151). Doric Neuroscience Studio software was used to modulate the LEDs and sample signals from the photoreceiver at 12 kHz, apply a 12 Hz low-pass filter, and decimate the signal to 120 Hz before writing the data to the hard drive. The start and end of every behavioral session were timestamped with TTL pulses from the VideoFreeze software and were recorded by photometry acquisition system to sync the photometry and behavioral data.

### Fiber photometry analysis

Fiber photometry data were analyzed using a custom python analysis pipeline. The fluorescence signals from 405-nm excitation and 465-nm excitation were downsampled to 10 Hz before calculating Δ*F*/*F*. Briefly, a linear regression model was fit to the 405 nm signal to predict the 465 nm signal. The predicted 465 nm signal was then used to normalize the actual 465 nm signal:

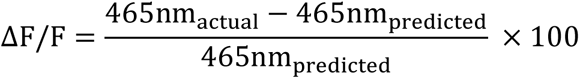

For analysis, individual TFC trials were extracted from the whole-session recording data, where each trial begins 20 s prior to CS onset and ends 100 s after the footshock. For each trial, Δ*F*/*F* values were z-scored using the 20 s baseline period prior to CS onset ((ΔF/F − μ_baseline_)/σ_baseline_).

Trial-averaged GCaMP responses were smoothed with loess regression for visualization purposes only; all statistical analyses were performed on the non-smoothed data. For statistical analysis, mean fluorescence values were calculated during the trace interval (“pre-shock”, 20-40 s after CS onset) and after the footshock (“post-shock”, 42-62 s after CS onset).

### Electrophysiological recordings

To improve SWRs detection, a custom-built microdrive^46^ holding 2-4 movable tetrodes (12.7um nickrome wire, Sandvik) allowed the vertical adjustment of each individual tetrode over the CA1 subfield. Tetrodes were lowered down once day and animals were recorded in an open field in the following day, until SWRs were visually identified; the tetrode lowering procedure lasted between 3-7 days. Local field potential (LFP) activity was recorded during TFC using a Cheetah data acquisition system (Digital Lynx 4SX, Neuralynx). LFP recordings were sampled at 2kHz, using a 0.3 - 500Hz bandpass filter. Recording data was synchronized with behavior using TTL pulses from the VideoFreeze software (Med Associates). To reduce electrical noise a lickometer switch (Med-Associates ENV-250S) was used to disconnect the shock generator from the grids during all periods when it was not in use. For anesthetized recordings, animals were injected (I.P.) with a ketamine (100 mg/kg), xylazine (15 mg/kg), acepromazine (2 mg/kg) mixture. Depth of anesthesia was monitored by tail pinch and respiratory rate. Animals were kept in their homecage during the entire recording. The experiment consisted of a baseline recording (5min duration) followed by an additional recording (5min duration) 10 minutes after the pilocarpine injection (10 mg/kg).

### Electrophysiological data analysis

Neuralynx data was converted to MATLAB format using a custom python script. The ripples were then analyzed in MATLAB using a custom-made routine. Raw signals were downsampled to 1kHz and bandpass filtered in the ripple band 130-200Hz. Putative SWR events were detect using the FindRipple function from FMAToolbox (https://fmatoolbox.sourceforge.net). SWR events were defined as those where the beginning/end thresholds exceeded 2 standard deviations and the peak exceeded 5 standard deviations. The minimum-maximum range for ripple duration was considered 20-100 ms, and a minimum of 30 ms for inter-ripple interval. Ripple frequency was classified between the ones that occurred during the baseline period of the TFC (2 minutes prior to the first tone presentation) or intertrial (2 minutes post-shock) periods.

### Statistical analysis

For analysis of the training and tone test behavioral data, freezing was measured during each trial epoch (session baseline, tone, trace, ITI) and averaged across trials for each animal. All behavioral data were analyzed using Two-Way Repeated Measures ANOVA followed by *post hoc* comparisons adjusted with the Sidak method when appropriate. For the context test session, freezing was computed across the entire session and analyzed using Welch’s unpaired t-test. For the fiber photometry a paired t-test was used to compare pre-shock and post-shock fluorescence within subjects. For the electrophysiological recordings a paired t-test was used to compare baseline with SWR frequency during intertrial intervals. A threshold of *p* < 0.05 was used to determine statistical significance. All data are shown as mean ± SEM. Statistical analyses were performed in python or GraphPad Prism version 10, and all figures were generated in python and BioRender.

### Histology

To verify viral expression and optical fiber location, mice were deeply anesthetized with isoflurane and transcardially perfused with cold phosphate buffered saline (1X PBS) followed by 4% paraformaldehyde (PFA) in 1X PBS. Brains were extracted and post-fixed with PFA overnight at room temperature. The following day 40 µm coronal sections were taken on a vibratome (Leica Biosystems) and stored in a cryoprotectant solution. Finally, slices containing the dorsal hippocampus were washed for 5 mins with 1X PBS three times before staining the slices for 10 minutes with DAPI (1:1,000, Life Technologies) and mounted on slides with Vectashield (Vector Labs). Images were acquired at 10x magnification on a fluorescence virtual slide microscope system (Olympus).

